# The genetics and genome-wide screening of regrowth loci, a key component of perennialism in *Zea diploperennis*

**DOI:** 10.1101/388256

**Authors:** Anjun Ma, Yinjie Qiu, Tajbir Raihan, Bimal Paudel, Subha Dahal, Yongbin Zhuang, Aravid Galla, Donald Auger, Yang Yen

## Abstract

Perennialism is common among the higher plants, yet little is known about its inheritance. To address this, six hybrids were made by reciprocally crossing perennial *Zea diploperennis* Iltis, Doebley & R. Guzman with inbred lines B73 and Mo17 and Rhee Flint, a heirloom variety, of *Z*. *mays* L. ssp. *mays*. All the F_1_ plants demonstrated several cycles of growth, flowering, senescence and regrowth into normal flowering plants, indicating a dominant effect of the Z. *diploperennis* alleles. The regrowability (i.e. the plants’ ability to regrow after senescence) was stably transmitted to progeny of the hybrids, so we focused on this trait. Segregation ratios in the F_2_ generations are consistent with the trait controlled by two dominant, complementary loci, but do not exclude the influence of other modifiers or environment. Genome-wide screening with genotyping-by-sequencing (GBS) indicated two major regrowth loci, *regrowth 1* and *regrowth 2*, were on chromosomes 2 and 7, respectively. These findings lay the foundation for further exploration of the molecular mechanism of regrowth in *Z. diploperennis*.

**Significance Statement:** This study contributes to the general understanding of inheritance of perennialism in the higher plants. Previous genetic studies of the perennialism in *Zea* have yielded contradictory results. We take a reductionist approach by specifically focusing on one trait, the plant’s ability to restart a new life cycle after senescence on the same body. While traits, such as rhizome formation, tillering and dormancy may be important to converting *Z. mays* to becoming truly perennial, understanding the conditions for regrowth after senescence will be substantial first step. Importantly, our data indicate that there is no major barrier to transferring this trait into maize or other grass crops for perennial crop development with proper technology, which enhances sustainability of grain crop production in an environmentally friendly way.

This manuscript was previously deposited as a preprint at http://dx.doi.org/10.1101/388256.

## Introduction

Perennialism is the phenomenon that a plant can live for more than two years; the ability of doing so is termed perenniality. Plants typically have a life cycle of growth, reproduction (sexual and/or vegetative) and senescence. Annuals and biennials have only one such cycle in their life, leaving behind seeds, bulbs, tubers, etc. to initiate another life cycle. Perennials maintain juvenile meristematic tissues capable of regrowth after senescence to start a new life cycle on the same body. How perennials do so remains as a mystery. Subterranean stems (such as rhizomes), polycarpy and tuberous roots are often cited as the means by which plants achieve perenniality. However, none of these traits is absolutely required by perennials. For instance, bamboos are essentially monocarpic perennial that regrow from rhizomes. Many perennial temperate grasses, such as switchgrass^(1)^, cordgrass^(2)^ and eastern gamagrass^(3)^, regrow from the crowns instead of rhizomes. On the other hand, some annual/biennial plants, such as radish (*Raphanus sativus*), grow tuberous roots.

Although perennialism is common among higher plants, the study of its genetics and molecular biology is sporadic. So far, the only published research in molecular mechanism of plant perennialism was conducted in Arabidopsis. Melzer et al. successfully mutated this annual herb to show some perennial habits, such as increased woody fiber in the stem, by down-regulating two flowering genes coding for MADS-box proteins, SUPPRESSOR OF OVEREXPRESSION OF CONSTANT 1 and FRUITFUL^(4)^. Unfortunately, this woody mutant was sterile, and no follow-up research was reported. Perennial-related genes and quantitative loci (QTL) have been reported in other species. Major QTL controlling rhizome development, regrowth and tiller number have been mapped on sorghum linkage groups C (chromosome 1) and D (chromosome 4)^(5, 6)^, which are homoeologous to regions of maize chromosomes 1, 4, 5 and 9, respectively^(7)^. Hu at al. mapped two dominant, complementary QTL *Rhz2* (*Rhizomatousness 2*) and *Rhz3* that control rhizome production on rice chromosomes 3 and 4 at the loci homoeologous to the sorghum QTL^(6)^. Tuberous roots in a wild perennial mungbean (*Vigna radiate* ssp. *sublobata*) are conditioned by two dominant, complementary genes^(8)^. However, after years of effort these perennialism-related genes have yet to be cloned from any of the species despite that mapping data and complete genomic sequences of rice and sorghum are readily available. Therefore, no further research has been reported about these perennialism-related loci/genes. Recently, Ryder and the associates reported a set of 98 expressed contigs in Johnsongrass (*S. halepense*) that are likely associated with rhizome development ^(9)^.

In the genus *Zea* L., most species, including maize (*Z*. *mays* ssp. *mays*), are annual. However, two closely related species, tetraploid *Z. perennis* [Hitchc.] Reeves and Mangelsdorf and diploid *Z. diploperennis* Iltis, Doebley & R. Guzman, are perennial. Perenniality of these two teosintes is manifested as regrowth after seed production and senescence, which includes developing juvenile basal axillary buds and rhizomes, under favorable environment. A fertile F_1_ hybrid between *Z. mays* and *Z. perennis* was made by Emerson^(10)^ in the 1920s, and *Z. mays*’ hybrids with *Z. diploperennis* were also obtained soon after the diploid perennial teosinte was discovered in the 1990s^(11)^. Evergreen stalks, bulbils (highly-condensed rhizomes), basal shoot development, stiff stalk and robust root system have all been cited as phenotypic features of perennialism in *Z. diploperennis*^(12-14)^. For example, evergreen stalks, which was proposed as a component of perennialism in Z*. diploperennis*^(12)^, appears to be linked to *sugary 1* on the short arm of chromosome 4^(15)^.

Conflicting conclusions have been reached in various studies on how perennialism is inherited in *Zea*. Shaver, who worked with tetraploid *Z. perennis*, proposed that a triple homozygous recessive genotype is needed for the perenniality in *Zea*^(16)^. In this model, *pe* (*perennialism*), interacting with *gt* (*grassy tillers*) and *id* (*indeterminate*), plays a key role in conferring totipotency to the basal axillary buds and rhizomes in the perennial teosintes^(16, 17)^. The nature of *pe* remains unknown and the *Z. perennis*-derived genotype from which *pe* was identified by Shaver^(16)^ was lost and never recovered despite decades of intensive efforts (Shaver, personal communication). Mangelsdorf and Dunn mapped *Pe*-d*, the maize allele of the *pe* homologue in *Z. diploperennis,* to the long arm of maize chromosome 4^(18)^. The *gt* gene (aka *gt1*), located on the short arm of maize chromosome 1, encodes a class I homeodomain leucine zipper that promotes lateral bud dormancy and suppresses elongation of lateral ear branches^(15)^. It appears that *gt1* depends on the activity of a major maize domestication gene, *teosinte branched 1* (*tb1*), and is inducible by shading^(19)^. The *id* gene (aka *id1*) alters maize’s ability to flower^(20)^. Both *tb1* and *id1* are located on the long arm of maize chromosome 1 and both encode transcription factors with zinc finger motifs^(19, 21)^. Singleton believed that *id1* inhibits plantlet generation at the upper nodes of a maize stalk^(20)^. Mangelsdorf and the associates proposed that one or two dominant genes control annual growth habit in their *Z. diploperennis*-popcorn hybrid^(22)^.

In contrast to the recessive inheritance model, Galinat proposed that perennialism in *Z. diploperennis* is at least partially controlled by two dominant complementary genes^(15)^. Also, Ting and Yu obtained three perennial F_1_ hybrids by pollinating three Chinese field corn varieties with *Z. diploperennis*^(23)^, which indicate that perennial factors are dominant. Unfortunately, there is no further report about these hybrids or their derivatives.

Westerbergh and Doebley regarded perennialism in *Z. diploperennis* as a quantitative trait and identified a total of 38 QTL for eight perennial-habit traits from a *Z. diploperennis* x *Z. mays* ssp. *parviglumis* (annual) mapping population^(12)^. Intriguingly, they did not identify any QTL that shows a singularly large effect. Murray and Jessup indicated that non-senescence and rhizomatousness are essential traits in their perennial maize breeding practice^(24)^.

Perennialism appears to be a complex trait, strongly influenced by genetic and environmental factors. A perennial plant in one environment usually cannot survive in another due to the lack of the required adaptability. For example, *Z. diploperennis*, which is perennial in the highlands of Mexico, cannot survive the harsh winter in the American Midwest. The various criteria for what constitutes perennialism in *Zea* may have contributed to contradictory observations. Traits such as rhizome formation, evergreen stalks, and dormancy are important adaptive features that support the viability of various perennial plants in various environments. In this study, we take a reductionist approach and specifically focus on a plant’s regrowability (i.e. the ability to maintain some juvenile meristematic tissues after each life cycle that can initiate a new life cycle). Although this trait by itself is insufficient for functional perenniality, it appears to be an essential component of perenniality in *Zea* L. Here we report the results of our genetic analysis and genome-wide screening of the regrowth trait with genotyping-by-sequencing (GBS) technology.

## Results and Discussion

### The production and growth of the hybrids

We made reciprocal crosses of *Z. diploperennis* (Zd, hereafter in a cross combination) with the following three maize lines: B73, Mo17 and Rhee Flint (RF, hereafter in a cross combination). B73 and Mo17 are inbred lines and Rhee Flint is an heirloom maize variety. The first F_1_ was made with Rhee Flint in a greenhouse. Rhee flint is small, fast-growing and usually has tillers, which affords serial plantings with an increased opportunity of a plant simultaneously flowering with *Z. diploperennis*. Because Rhee Flint is an open-pollinated variety, later F_1_s were made with B73 and Mo17 to facilitate molecular analysis. All the F_1_ plants are fertile and completed multiple cycles of growth, reproduction and senescence (Fig. 1; Supplementary Fig. S1). Regrowth (as opposed to accidental replanting from seed) of F_1_ plants was insured by inspection that new shoots were attached to the base of the F_1_ and confirmed by the heterozygosity of polymorphic PCR markers (examples shown in Supplementary Fig. S2). Regrowth of these F_1_s originates from basal axillary buds after stem senescence in all the crosses (Figs. 1D, 1E, 1F).

**Figure 1.**
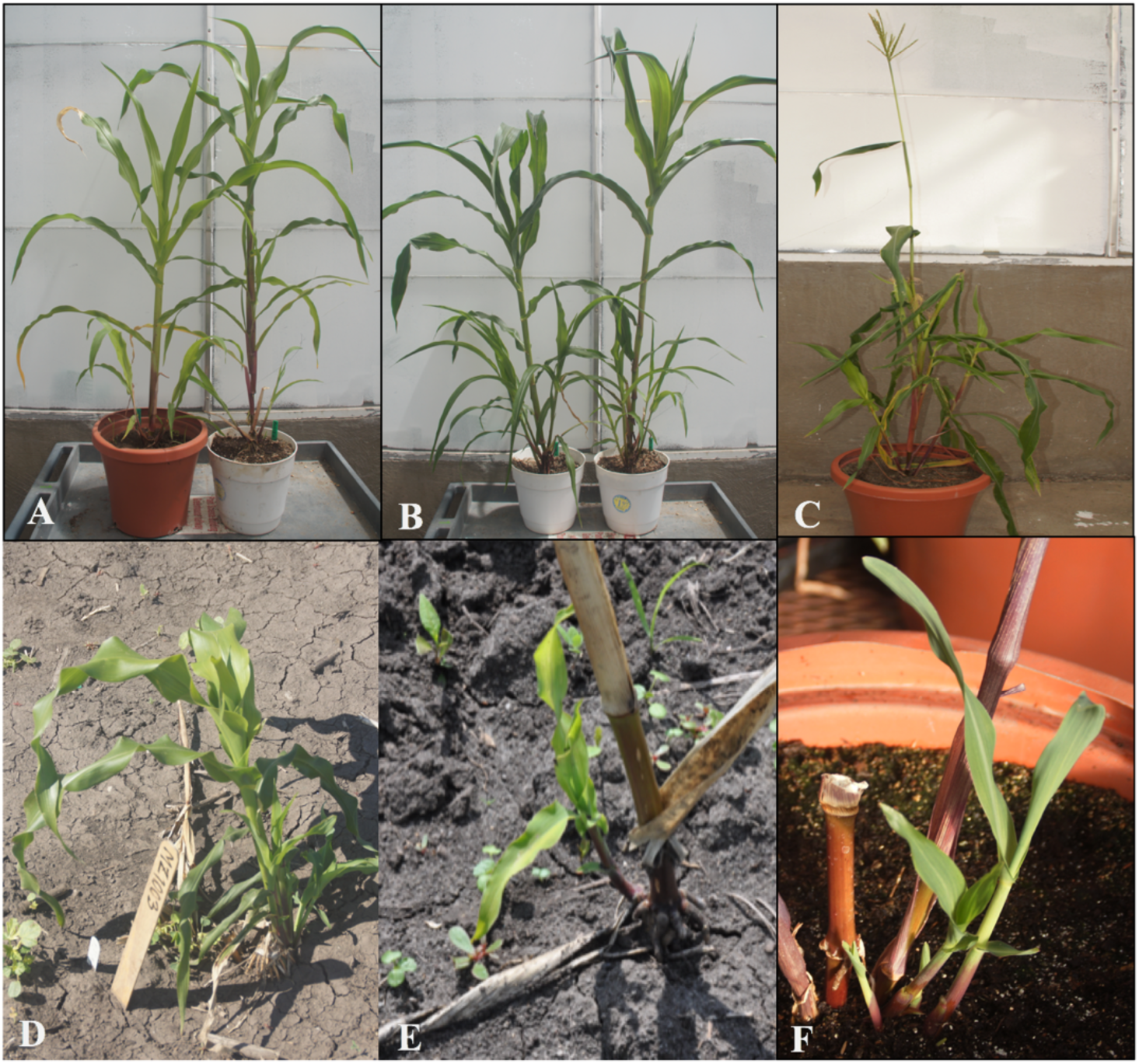
Photos of *Zea mays* and *Z. diploperennis* (Zd) F_1_ plants. **A**: reciprocal Mo17-Zd (right) and Zd-Mo17 (left) F_1_ plants; **B**: reciprocal B73-Zd (right) and Zd-B73 (left) F_1_ plants; **C**: RF-Zd F_1_ plant; **D**: regrowth of a Mo17-Zd F_1_ plant; **E**: regrowth of a B73-Zd F_1_ plant; and **F**: regrowth of a RF-Zd F_1_ plant. B73, Mo17 and RF represent, respectively, inbred lines B73 and Mo17 and cultivar Rhee Flint of *Z. mays*.

Some of the basal regrowth immediately developed into a female (Fig. 2A) or a male (Fig. 2B) inflorescence, or a forest of them (Fig. 2D). These F_1_ plants with abnormal regrowth most often can later undergo normal regrowth in an alternative environment, such as being moved from the greenhouse to the field, etc., which suggests a strong environmental influence. No such abnormal growth has been seen in advanced generations. Sometimes, plantlets also can develop at the upper nodes of some hybrid plants when B13 and Mo17 were used as the parent (Fig. 2C). The plantlets developed at the upper nodes, however, can only survive if transplanted into soil. This indicates that the senescent stalks do not function to provide the necessary nutrients to the plantlets.

**Figure 2.**
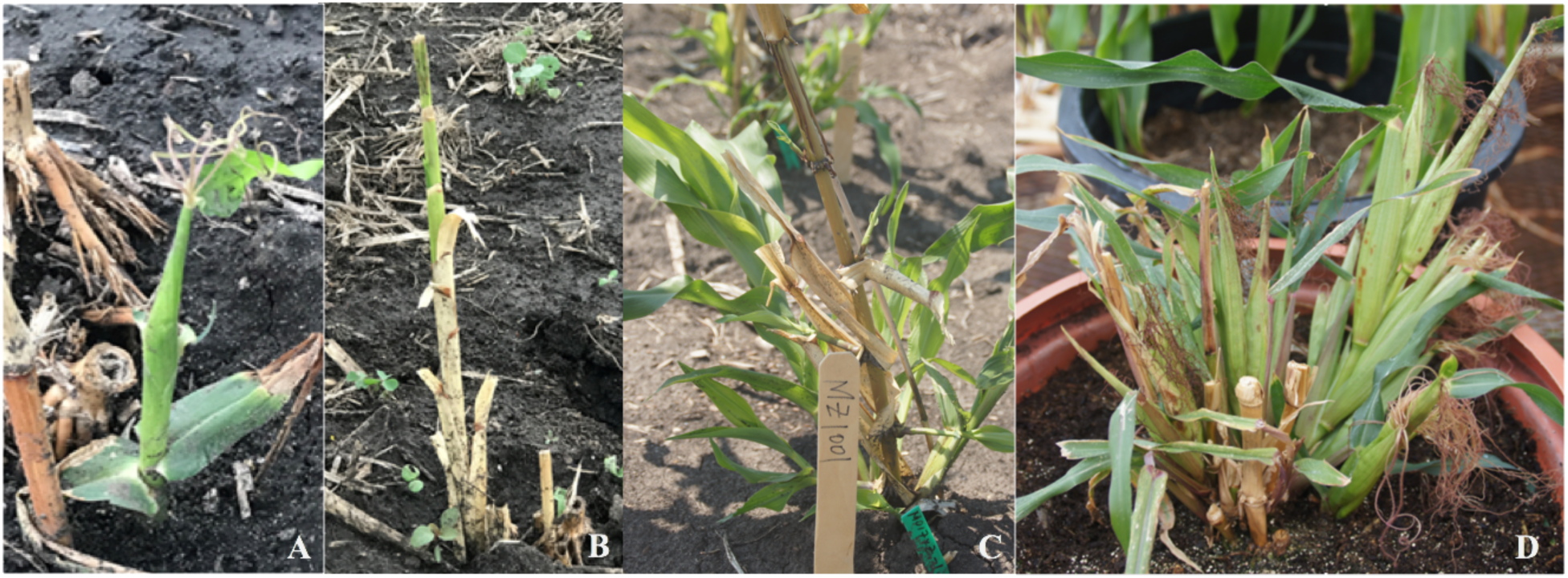
Photos of abnormal F_1_ plants of crosses of *Zea diploperennis* with *Z. mays* inbred lines B73 (A & B) and Mo17 (C) or cv. Rhee Flint (D).

Because the F_1_ plants and their perennial derivatives are not winter hardy, the regeneration cycles were alternated between the greenhouse and the field (Supplementary Figs. S1 & S3). Interestingly, the ears and kernels of the F_1_s of the six crosses all were more teosinte-like (i.e. two rows of oppositely positioned spikelets with paired kernels encased by woody rachides and glumes) when grown in greenhouse but were more maize-like (i.e. multiple rows of naked kernels with short soft glumes and rachides around a silica-filled soft core) when grown in the field (Fig. 3). In the F_2_ and higher generations, ear morphology segregated even under greenhouse condition (Fig. 3). These observations suggest that environmental factors are important in the preferential expression of the teosinte or the maize alleles of the genes influencing ear morphogenesis in the hybrids. These observations also indicate that it should be possible to breed regrowable maize with maize-like ears and kernels.

**Figure 3.**
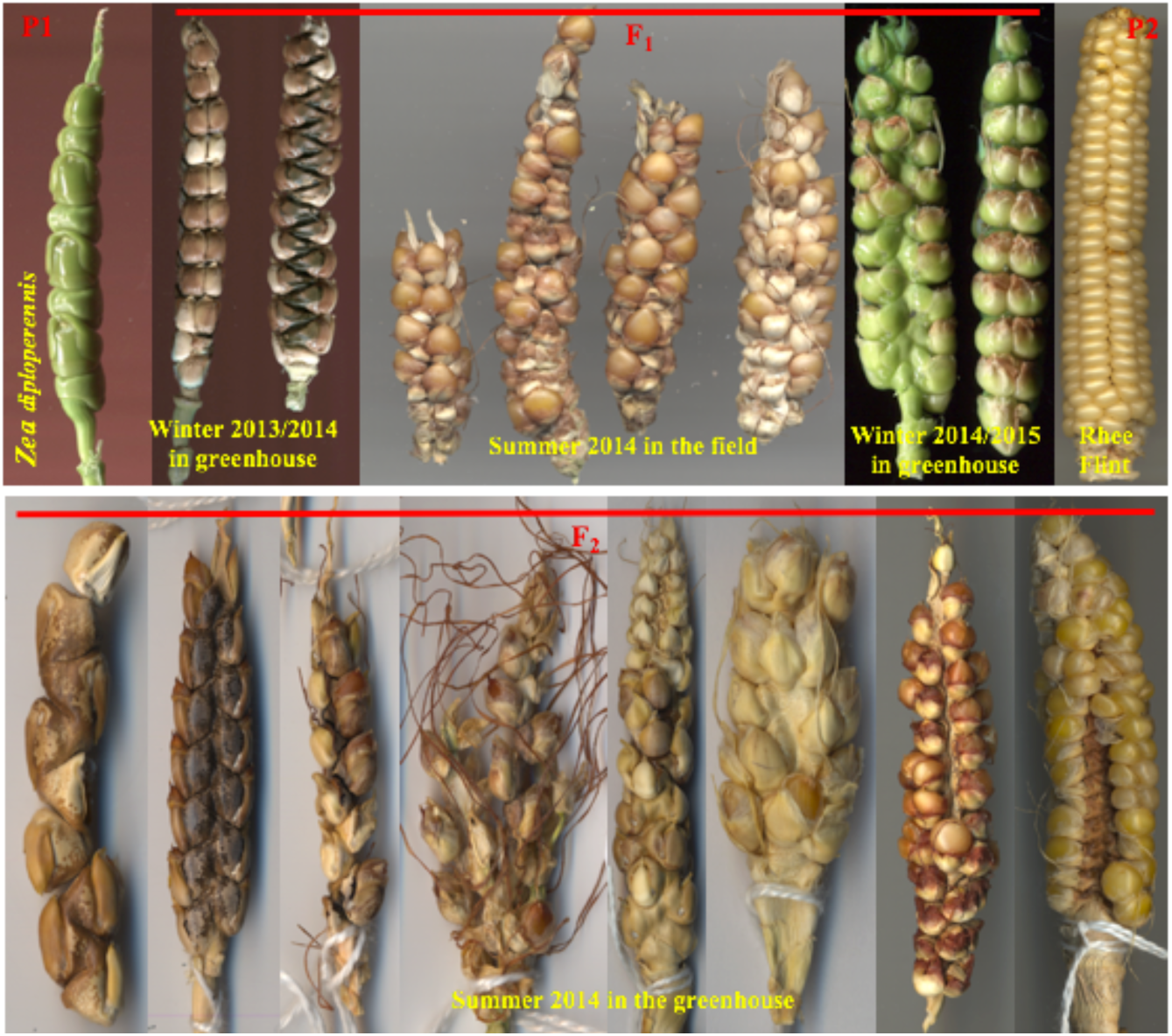
Photos of the ears produced from a *Zea mays* cv Rhee Flint x *Z. diploperennis* F_1_ plant in different seasons (the upper panel) and from F_2_ in summer 2014 in greenhouse (the lower panel).

The contrast between our observations and those of some previous reports is remarkable. While we focus on a single trait, regrowth after senescence, previous studies were interested in perennialism generally using varying criteria. Conclusions that perennialism in *Zea* is recessive might have resulted from the hypothesis that traits such as tiller number at tasseling (TNT) or rhizome development are indispensable components of perennialism in *Zea*. Indeed, other studies have used rhizome development as an indicator of perennialism in *Zea*^(11, 16, 17, 22, 25)^ and we have not observed rhizomes in any of our F_1_s or the derived plants. When regrowth occurs, it is always from an axillary bud. Indeed, it is also our observation that the regrowth of *Z. diploperennis* is mainly from basal axillary buds, and only occasionally from rhizomes. The *Z. perennis* - 4X maize F_1_s made by R. A. Emerson also were all “weekly perennial” under the environmental conditions with few or no rhizomes^(10)^.

Other possible explanations for contrasting results is that the perennial teosinte plants used in those studies were polymorphic for one or more regrowth genes, that the experimental environments were unfavorable for regrowth to happen, or that some plants needed more time to break up their dormancy. Shaver^(16)^ and Camara-Hernandez and Mangelsdorf^(25)^ observed that some of their F_1_ plants eventually regrew from basal axillary buds after a period of dormancy. Indeed, some of our F_2_ plants need about two months of dormancy before regrowth. This observation reinforces the view that even regrowth is a complex trait that is modified by genetics and environment.

TNT has been associated with perennialism in several studies^(14, 17, 26, 27)^, so we investigated the relationship of TNT with regrowth in the Zd-RF F_2_s. One-way ANOVA of TNT by regrowth (Supplementary Table S1), however, revealed no significant difference of TNT (*F* = 0.897, *p* = 0.353) between the regrowth and the non-regrowth F_2_s. Indeed, we observed regrowth from several single-stalked hybrid derivatives (Fig. 4A) and non-regrowth of some multi-stalked plants (Fig. 4B). These results suggest that TNT is not essential to regrowth in *Zea*.

**Figure 4.**
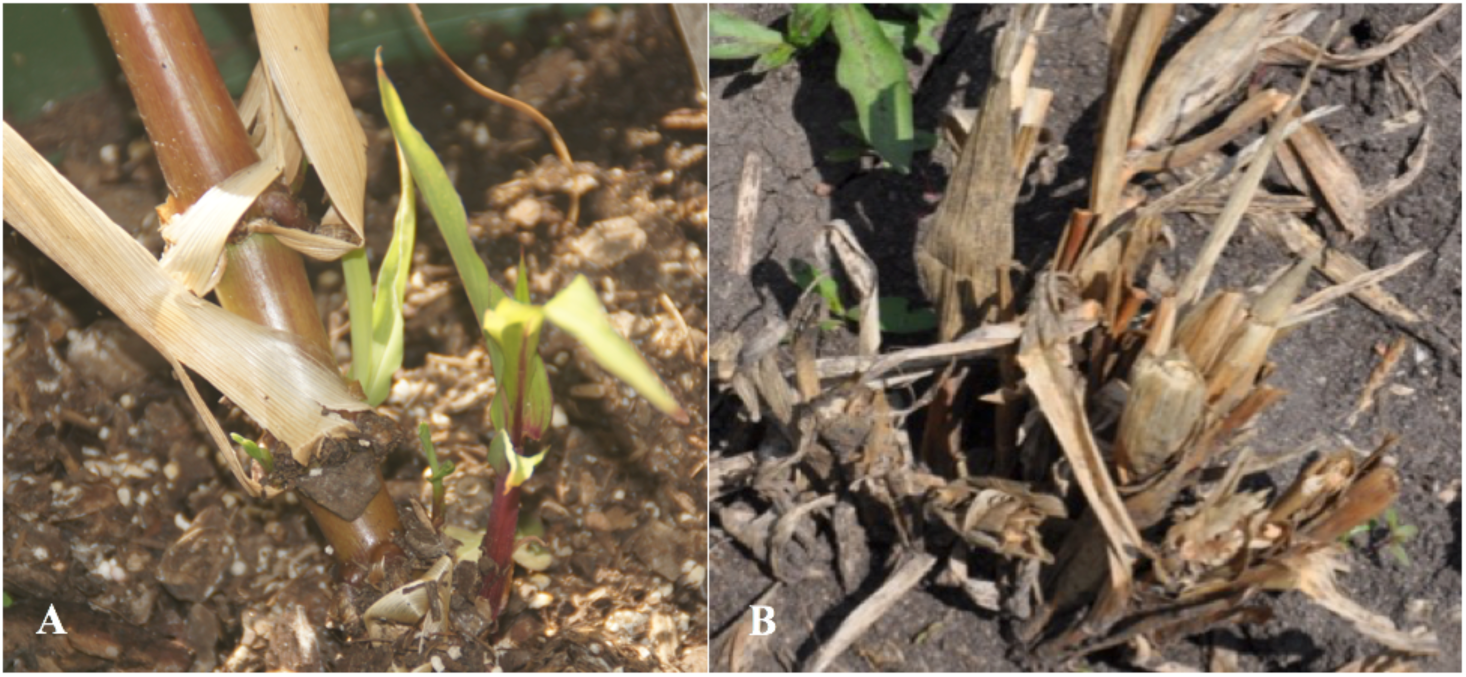
Photos of *Zea mays* Mo17-*Z*. *diploperennis* F_2_ plants, showing regrowth from the basal node of a single-stalked plant (A) or non-regrowth from a multi-stalked plant (B).

### The genetics of regrowth

All our F_1_ plants regrew and underwent several life cycles alternating between the greenhouse and the field. This indicates that, with our materials and in our environment, regrowth is a dominant trait. Although this contrasts with some reports, regrowable F_1_ hybrids of maize with perennial teosintes were previously obtained by Emerson^9^, Shaver^(16)^, Galinat^(15)^, Ting and Yu^(24)^ and Camara-Hernandez and Mangelsdorf^(25)^. Brewbaker suggested cytoplasm may contribute to perennialism^(28)^, but our reciprocal F_1_s performed similarly.

To analyze the genetics of regrowth further, 134 B73-Zd F_2_s (derived from several F_1_ plants where B73 was the female) and 159 Zd-RF F_2_s (derived from a single F_1_ plant where Zd was the female) were tested. Among the 134 B73-Zd F_2_s, 81 regrew and 53 did not (Table 1). Similarly, among the 159 Zd-RF F_2_s, 90 regrew after senescence and 69 did not (Table 2). One B73-Zd F_3_ population (Supplementary Table S1) and three Zd-RF F_3_ populations (Table 2), each of which was derived from a single regrowth F_2_ plant, were also evaluated for their regrowth.

**Table 1.**
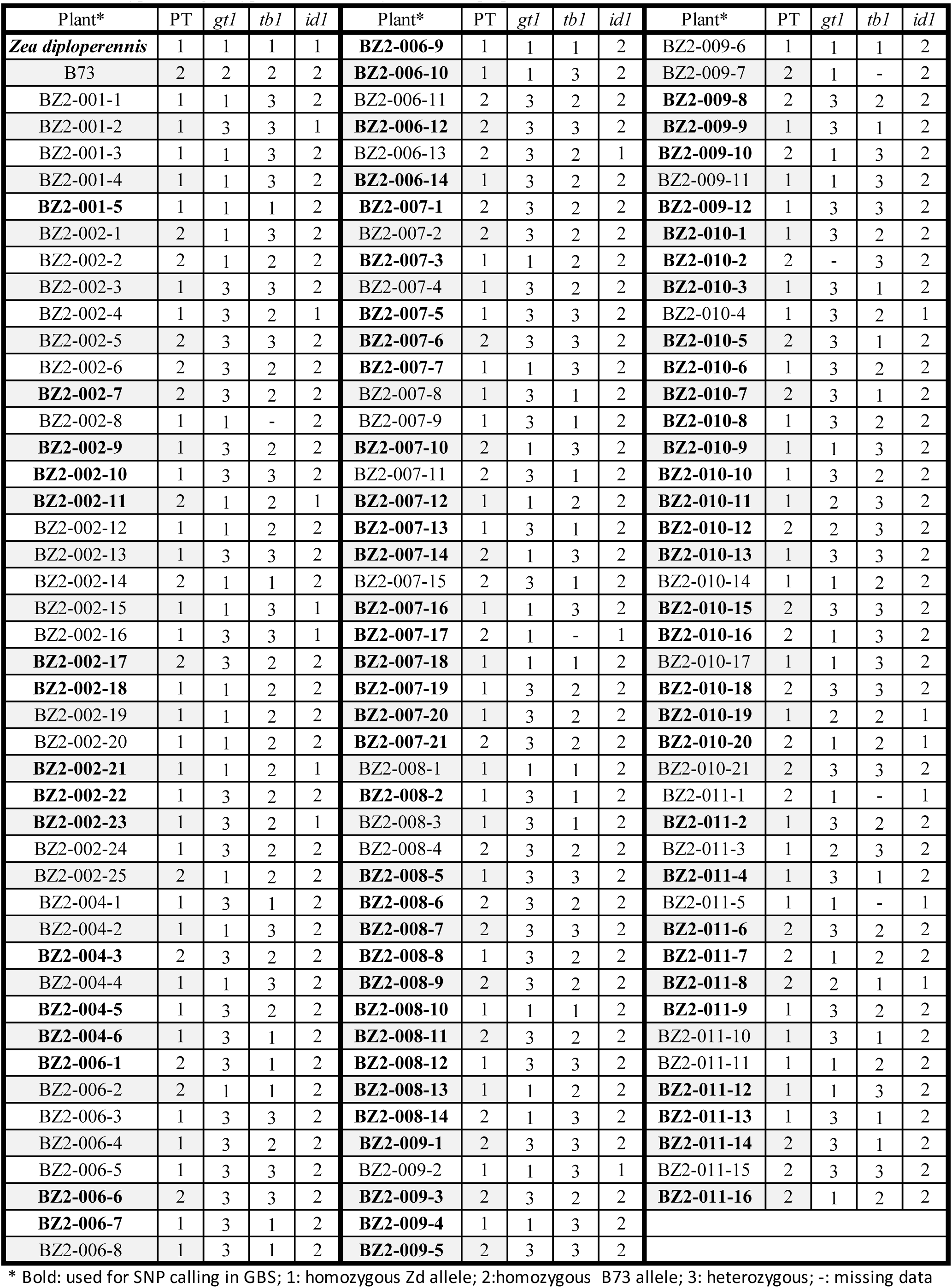
The regrowth (R) and the non-regrowth (NR) phenotypes and the marker genotypes of *Zea diploperennis* (Zd), *Z. mays* B73 and their 134 F_2_ plants*.

**Table 2.** Segregation of regrowth among the *Zea diploperennis*-*Z. mays* cv Rhee Flint F_2_s and F_3_s.

| Plant | PT | Plant | PT | Plant | PT | Plant | PT | Plant | PT | Plant | PT | Plant | PT |
| --- | --- | --- | --- | --- | --- | --- | --- | --- | --- | --- | --- | --- | --- |
| ZR2-001-1 | 1 | ZR2-001-33 | 2 | ZR2-001-69 | 0 | ZR2-001-102 | 1 | ZR2-001-132 | 2 | ZR2-001-162 | 2 | ZR3-005-7 | 1 |
| ZR2-001-2 | 2 | ZR2-001-34 | 1 | ZR2-001-71 | 1 | ZR2-001-103 | 1 | ZR2-001-133 | 1 | ZR2-001-163 | 1 | ZR3-005-8 | 1 |
| ZR2-001-3 | 1 | ZR2-001-35 | 2 | ZR2-001-72 | 0 | ZR2-001-104 | 1 | ZR2-001-134 | 1 | ZR2-001-164 | 1 | ZR3-005-9 | 1 |
| ZR2-001-4 | 2 | ZR2-001-36 | 2 | ZR2-001-73 | 1 | ZR2-001-105 | 1 | ZR2-001-135 | 2 | ZR2-001-165 | 2 | ZR3-005-10 | 1 |
| ZR2-001-5 | 1 | ZR2-001-37 | 2 | ZR2-001-74 | 1 | ZR2-001-106 | 1 | ZR2-001-136 | 2 | ZR2-001-166 | 1 | ZR3-005-11 | 2 |
| ZR2-001-6 | 1 | ZR2-001-38 | 2 | ZR2-001-75 | 1 | ZR2-001-107 | 2 | ZR2-001-137 | 1 | ZR2-001-167 | 2 | ZR3-005-12 | 2 |
| ZR2-001-7 | 2 | ZR2-001-39 | 1 | ZR2-001-77 | 0 | ZR2-001-108 | 2 | ZR2-001-138 | 1 | ZR2-001-168 | 2 | ZR3-005-13 | 1 |
| ZR2-001-9 | 1 | ZR2-001-40 | 2 | ZR2-001-78 | 0 | ZR2-001-109 | 2 | ZR2-001-139 | 1 | ZR2-001-169 | 2 | ZR3-005-14 | 2 |
| ZR2-001-10 | 2 | ZR2-001-42 | 2 | ZR2-001-79 | 1 | ZR2-001-110 | 1 | ZR2-001-140 | 2 | ZR2-001-171 | 1 | ZR3-005-15 | 2 |
| ZR2-001-11 | 1 | ZR2-001-43 | 1 | ZR2-001-80 | 1 | ZR2-001-111 | 2 | ZR2-001-141 | 1 | ZR3-003-1 | 1 | ZR3-005-16 | 2 |
| ZR2-001-12 | 1 | ZR2-001-44 | 1 | ZR2-001-81 | 1 | ZR2-001-112 | 1 | ZR2-001-142 | 1 | ZR3-003-2 | 1 | ZR3-009-1 | 1 |
| ZR2-001-13 | 2 | ZR2-001-45 | 1 | ZR2-001-82 | 0 | ZR2-001-113 | 2 | ZR2-001-143 | 1 | ZR3-003-3 | 1 | ZR3-009-2 | 1 |
| ZR2-001-14 | 2 | ZR2-001-47 | 2 | ZR2-001-83 | 0 | ZR2-001-114 | 2 | ZR2-001-144 | 2 | ZR3-003-4 | 2 | ZR3-009-3 | 1 |
| ZR2-001-15 | 1 | ZR2-001-48 | 2 | ZR2-001-84 | 1 | ZR2-001-115 | 2 | ZR2-001-145 | 1 | ZR3-003-6 | 1 | ZR3-009-4 | 1 |
| ZR2-001-16 | 2 | ZR2-001-49 | 1 | ZR2-001-85 | 1 | ZR2-001-116 | 1 | ZR2-001-146 | 1 | ZR3-003-7 | 1 | ZR3-009-5 | 1 |
| ZR2-001-17 | 1 | ZR2-001-51 | 2 | ZR2-001-86 | 1 | ZR2-001-117 | 1 | ZR2-001-147 | 1 | ZR3-003-8 | 1 | ZR3-009-6 | 2 |
| ZR2-001-18 | 2 | ZR2-001-53 | 2 | ZR2-001-87 | 0 | ZR2-001-118 | 2 | ZR2-001-148 | 1 | ZR3-003-9 | 1 | ZR3-009-7 | 1 |
| ZR2-001-19 | 2 | ZR2-001-54 | 1 | ZR2-001-88 | 0 | ZR2-001-119 | 1 | ZR2-001-149 | 1 | ZR3-003-10 | 1 | ZR3-009-8 | 1 |
| ZR2-001-20 | 1 | ZR2-001-55 | 1 | ZR2-001-89 | 0 | ZR2-001-120 | 1 | ZR2-001-150 | 1 | ZR3-003-11 | 2 | ZR3-009-9 | 1 |
| ZR2-001-21 | 2 | ZR2-001-56 | 1 | ZR2-001-90 | 1 | ZR2-001-121 | 2 | ZR2-001-151 | 1 | ZR3-003-12 | 1 | ZR3-009-10 | 1 |
| ZR2-001-22 | 2 | ZR2-001-57 | 1 | ZR2-001-91 | 1 | ZR2-001-122 | 1 | ZR2-001-152 | 2 | ZR3-003-13 | 1 | ZR3-009-11 | 1 |
| ZR2-001-23 | 1 | ZR2-001-58 | 2 | ZR2-001-92 | 1 | ZR2-001-123 | 2 | ZR2-001-153 | 1 | ZR3-003-14 | 1 | ZR3-009-12 | 1 |
| ZR2-001-24 | 1 | ZR2-001-59 | 1 | ZR2-001-93 | 1 | ZR2-001-124 | 1 | ZR2-001-154 | 1 | ZR3-003-15 | 1 | ZR3-009-13 | 1 |
| ZR2-001-25 | 2 | ZR2-001-60 | 1 | ZR2-001-94 | 0 | ZR2-001-125 | 1 | ZR2-001-155 | 2 | ZR3-003-16 | 2 | ZR3-009-14 | 2 |
| ZR2-001-26 | 1 | ZR2-001-62 | 1 | ZR2-001-95 | 0 | ZR2-001-126 | 2 | ZR2-001-156 | 1 | ZR3-005-1 | 1 | ZR3-009-15 | 2 |
| ZR2-001-27 | 2 | ZR2-001-63 | 1 | ZR2-001-97 | 1 | ZR2-001-127 | 2 | ZR2-001-157 | 2 | ZR3-005-2 | 1 | ZR3-009-16 | 1 |
| ZR2-001-28 | 2 | ZR2-001-64 | 1 | ZR2-001-98 | 1 | ZR2-001-128 | 2 | ZR2-001-158 | 2 | ZR3-005-3 | 2 | PT: phenotype<br>1: regrowth<br>2: non-regrowth |  |
| ZR2-001-30 | 1 | ZR2-001-65 | 1 | ZR2-001-99 | 1 | ZR2-001-129 | 1 | ZR2-001-159 | 2 | ZR3-005-4 | 2 |  |  |
| ZR2-001-31 | 1 | ZR2-001-67 | 2 | ZR2-001-100 | 1 | ZR2-001-130 | 2 | ZR2-001-160 | 1 | ZR3-005-5 | 1 |  |  |
| ZR20-01-32 | 1 | ZR2-001-68 | 2 | ZR2-001-101 | 1 | ZR2-001-131 | 2 | ZR2-001-161 | 1 | ZR3-005-6 | 1 |  |  |

A chi square (*χ2*) goodness-of-fit test suggests that both of the F_2_ populations and one Zd-RF F_3_ population (Zd-RF F_3_-5) we tested best fit a 9:7 regrowth to non-regrowth ratio (Table 3), and the B73-Zd F_3_ population and the remaining two Zd-RF F_3_ populations best fit a 3:1 ratio (Table 3). The simplest model that explains these results is that regrowth in the F_1_s and their derivatives is mainly controlled by two dominant, complementary *regrowth* (*reg*) loci. A two dominant, complementary gene model parallels what has been found in other species, such as rice (*Oryza sativa*)^(6)^, Johsongrass (*Sorghum halepense*)^(5, 6, 29)^, basin wildrye (*Leymus cinereus*)^(30)^ and wild mungbean (*Vigna radiate* ssp. *sublobata*)^(8)^, for perennialism-related traits.

**Table 3.** Results of the χ2 goodness-of-fit tests of three genetic models.

| Generations | Observed | | | No. dominant genes (the expected R to NR ratio) and $P(\chi^2)^*$ | | |
| --- | --- | --- | --- | --- | --- | --- |
|  | Total | R | NR | 1 (3:1) | 2 (9:7) | 3 (27:37) |
| B73-Zd F<sub>2</sub> | 134 | 81 | 53 | 0.0001 | **0.2964** | 0.0001 |
| B73-Zd F<sub>3</sub> | 72 | 52 | 20 | **0.5862** | 0.0063 | 0.0001 |
| Zd-RF F<sub>2</sub> | 160 | 92 | 68 | 0.0001 | **0.7499** | 0.0001 |
| Zd-RF F<sub>3</sub>-3 | 15 | 12 | 3 | **0.6547** | 0.0639 | 0.3000 |
| Zd-RF F<sub>3</sub>-5 | 16 | 9 | 7 | 0.0833 | **1.0000** | 0.2547 |
| Zd-RF F<sub>3</sub>-9 | 16 | 13 | 3 | **0.5637** | 0.0438 | 0.0016 |

\*: the best fit models are in bold.

The Zd-RF F_1_ was also backcrossed to each parental line. All plants from the F_1_-to-Zd backcross regrew, showing dominant effect of the Zd alleles. However, only one of the 20 plants from the F_1_-to-RF backcross showed regrowth. Therefore, alternative models, such as one or three dominant complementary genes, two major genes with a few minor modifiers, or that regrowth is a complex trait controlled by many QTL, are not eliminated by this genetic analysis, but are less probable.

The number of regrowth plants observed in any generation might be understated, because some plants initially recorded as non-regrowth eventually regrew after about two months of dormancy. Therefore, some plants recorded as non-regrowth and discarded to open up greenhouse space may have possessed the ability to regrow. Furthermore, transplanting from the field to the greenhouse and *vise versa* is very stressful so that some regrowable plants may have been killed this way.

The F_2_ and F_3_ plants afforded an opportunity to investigate whether some factors previously implicated in perennialism may contribute to the regrowth trait. The rice rhizomatousness gene *Rhz2* has been mapped to rice chromosomes 3^(6)^ and sorghum chromosome 1^(5, 6, 29)^, which are both homoeologous to parts of maize chromosome 1^(7)^. Additionally, *gt1* and *id1*, which have been implicated with perenniality in *Zea*^(8)^,and *tb1*, which controls *gt1*^(19)^, are all on chromosome 1 in *Zea*^(19, 21)^. Therefore, we investigated the allele compositions of these three genes in the B73-Zd F_2_s (Table 1), and 26 Zd-RF F_2_ plants and the three Zd-RF F_3_ populations (Supplementary Table S2), and assayed their association with regrowth. Of the 134 regrowth hybrid derivatives we examined, 5, 33 and 115 were homozygous for the maize *gt1*, *tb1* or *id1* alleles, respectively (Table 1 and Supplementary Table S2). Zd-RF F_3_ family Zd-RF F_3_-5 is homozygous for the *gt1* allele of *Z. diploperennis* (Supplementary Table S2) but segregates approximately 9:7 for regrowth and non-regrowth (Table 2). Therefore, our results are inconsistent with the model of Shaver^(16)^, and show that *gt1* and *id1* do not control regrowth in our F_1_s and their derivatives. *Z*. *diploperennis*’s *gt1* allele may be helpful to regrowth because the majority of the plants that regrew had at least one copy, but it is not indispensable because some plants regrew without it.

Interestingly, we observed no heterozygosity for *id1* and very low heterozygosity for *tb1* in all the hybrid derivatives that were examined, regardless of regrowth (Tables 1; Supplementary Table S2). Of 134 B73-Zd F_2_ plants, only 16 had the *Z. diploperennis id1* allele (Table 1). Similar phenomena were observed in the derivatives of the Zd-RF cross (Supplementary Table S2). It seems that the maize chromosome fragment that carries *id1* was preferentially transmitted to the hybrid derivatives. Excess homozygosity of the maize *id1* allele indicates some sort of selection. It could be that a deficiency or other rearrangement adjacent to the teosinte *id1* allele causes it not to transmit efficiently, or it could be that the teosinte *id1* or tightly linked allele causes the plant not to grow well or flower in our experimental conditions.

### Identifying regrowth loci with genotyping-by-sequencing assay

A genome-wide mining of single nucleotide polymorphisms (SNPs) was conducted in a randomly selected sub-population of 94 (55 regrowth and 39 non-regrowth) B73-Zd F_2_ plants with GBS technology (Table 1). We conducted the GBS assays to identify QTL for the regrowth trait. To prepare for these assays, a total of 2,204,834 (85.14%) Illumina sequencing tags that passed routine quality control filtrations were aligned with the B73_v4 reference genome. A total of 714,158 SNPs, covering all ten chromosomes with an average of 71,416 SNPs per chromosome, were then called from 83 (46 regrowth and 37 non-regrowth, labeled in bold in Table 1) of the 94 F_2_ plants using TASSEL 3 pipeline^(31, 32)^. SNP-calling for the excluded 11 plants probably failed due to an error in barcode addition before sequencing. These SNPs were first subjected to a two-step filtration in TASSEL 3 (MergeDuplicateSNPsPlugin and GBSHapMapFiltersPlugin) to remove SNPs with poor qualities (see the Materials & Methods section and Supplementary Table S3 for the details), which resulted in 37,925 SNPs (Table 4, 1^st^ filtration). The 37,925 SNPs were then manually filtered with high missing data rate > 20%, which resulted in 10,432 remaining SNPs among all ten chromosomes (Table 4, 2^nd^ filtration).

**Table 4.** The actions taken and the numbers of SNPs revealed in each filtration step of SNP analysis of *Zea mays* B13 - *Z. diploperennis* F_2_ plants.

| **Chr** | **Raw SNP number** | **1<sup>st</sup> filter** | **2<sup>nd</sup> filter** | **3<sup>rd</sup> filter** | **4<sup>th</sup> filter** |
| --- | --- | --- | --- | --- | --- |
| **1** | 109,543 | 5,751 | 1,628 | 82 | 51 |
| **2** | 85,283 | 4,966 | 1,476 | 120 | 77 |
| **3** | 81,625 | 4,708 | 1,200 | 120 | 75 |
| **4** | 75,832 | 3,376 | 942 | 112 | 82 |
| **5** | 77,314 | 4,409 | 1,197 | 198 | 111 |
| **6** | 58,195 | 2,938 | 761 | 87 | 49 |
| **7** | 62,280 | 3,108 | 877 | 144 | 98 |
| **8** | 57,748 | 3,210 | 877 | 20 | 16 |
| **9** | 57,231 | 2,982 | 741 | 29 | 16 |
| **10** | 49,107 | 2,477 | 732 | 34 | 22 |
| **Total** | **714,158** | **37,925** | **10,431** | **946** | **597** |

To explore which chromosomal regions may control the regrowth phenotype, the 10,432 SNPs from the 2^nd^ filtration step were pooled for QTL analysis using R/qtl (version 1.42-8) (Supplementary Table S4). The result is shown in Figure 5A. A total of 126 SNPs (104 real sites and 22 simulated sites) showed LOD scores higher than 3.00. A permutation test of 1,000 with the *p-*value of 0.05 resulted in a significant LOD score of 5.23 (Fig. 5A). This significance threshold revealed two major QTL with one at 33,041,409 bp (the nucleotide position in the B73_v4 reference genome sequence) on chromosome 2 with a LOD score of 5.46 and one at 4,284,633 bp on chromosome 7 with a LOD score of 5.53, respectively.

**Figure 5.**
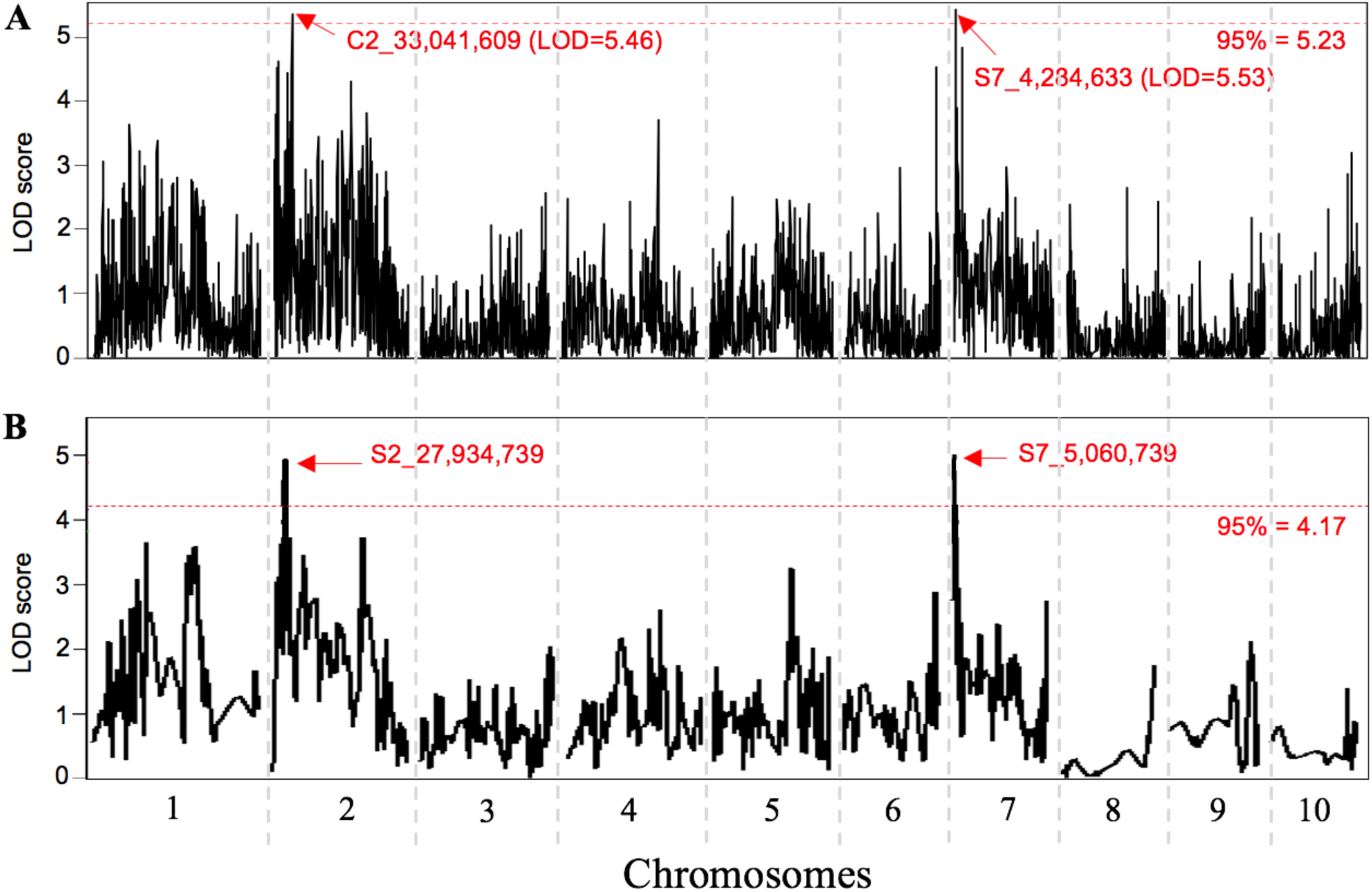
Graphics showing LOD scores of the QTL mapping the B73-Zd F2 population without (A) or with (B) chi-square imputation. The 95% threshold lines (the parallel red dash lines) were calculated with 1,000 permutation. Significant QTL/loci are indicated by the location of the peak SNPs of the loci.

To test if the two strongest QTL correspond to two dominant and complementary loci suggested by the genetic analysis, we applied a *χ2* test with 9:7 allele segregation ratio model to the 10,431 SNPs to investigate if the observed and the expected genotypes are significantly different (*p* ≤ 0.05). This is based on our hypothesis that, if a SNP is associated with a *reg* locus, the teosinte allele of the SNP should be carried by all the regrowable F_2_s but only by three sevenths non-regrowable F_2_s. This step kept 946 SNPs (Table 4, 3^rd^ filtration). Finally, to simplify the mapping effort, the 946 SNPs were filtered once more by collapsing immediately neighboring SNPs that share the same haplotypes into one cluster. This step resulted in 597 SNP cluster with each being represented by the first SNP in the cluster (Table 4, 4^th^ filtration). Locus mapping was then conducted with a threshold of LOD^95%^ = 4.17. This analysis revealed two significant loci with one on chromosome 2 in the interval from 24,244,192 bp to 28,975,747 bp with the peak at 27,934,739 bp and another on chromosome 7 in the interval from 2,862,253 bp to 6,681,861 bp with the peak at 5,060,739 bp (Fig. 5B). These two loci were mapped closely to the two major QTL indicated by the mapping without imputation. These results are consistent with the two-factor model, which warrants further investigation. To that end we are naming the factor underlying the chromosome 2 QTL *regrowth 1* (*reg1*) and the factor underlying the chromosome 7 QTL as *regrowth 2* (*reg2*).

Our LOD analysis located two minor peaks on chromosome 1 that are associated with regrowth (Fig. 5B). We wanted to know if these two loci are related to *gt1* and *id1*, respectively. The SNPs at the peak of these loci are at 82,273,951 bp and 177,235,112 bp, far away from *id1* (around 243,201,405 bp) and *gt1* (around 23,625,801 bp) (Fig. 6). These observations further indicate that *id1* and *gt1* are not related to regrowth. Also, previous studies reported that *Z. diploperennis* carried perennialism-related *Pe*-d* and an evergreen gene on chromosome 4^(15, 19)^. However, our data could not support these observations since no SNP on chromosome 4 is significantly associated with regrowth (Figs. 5 and 6).

**Figure 6.**
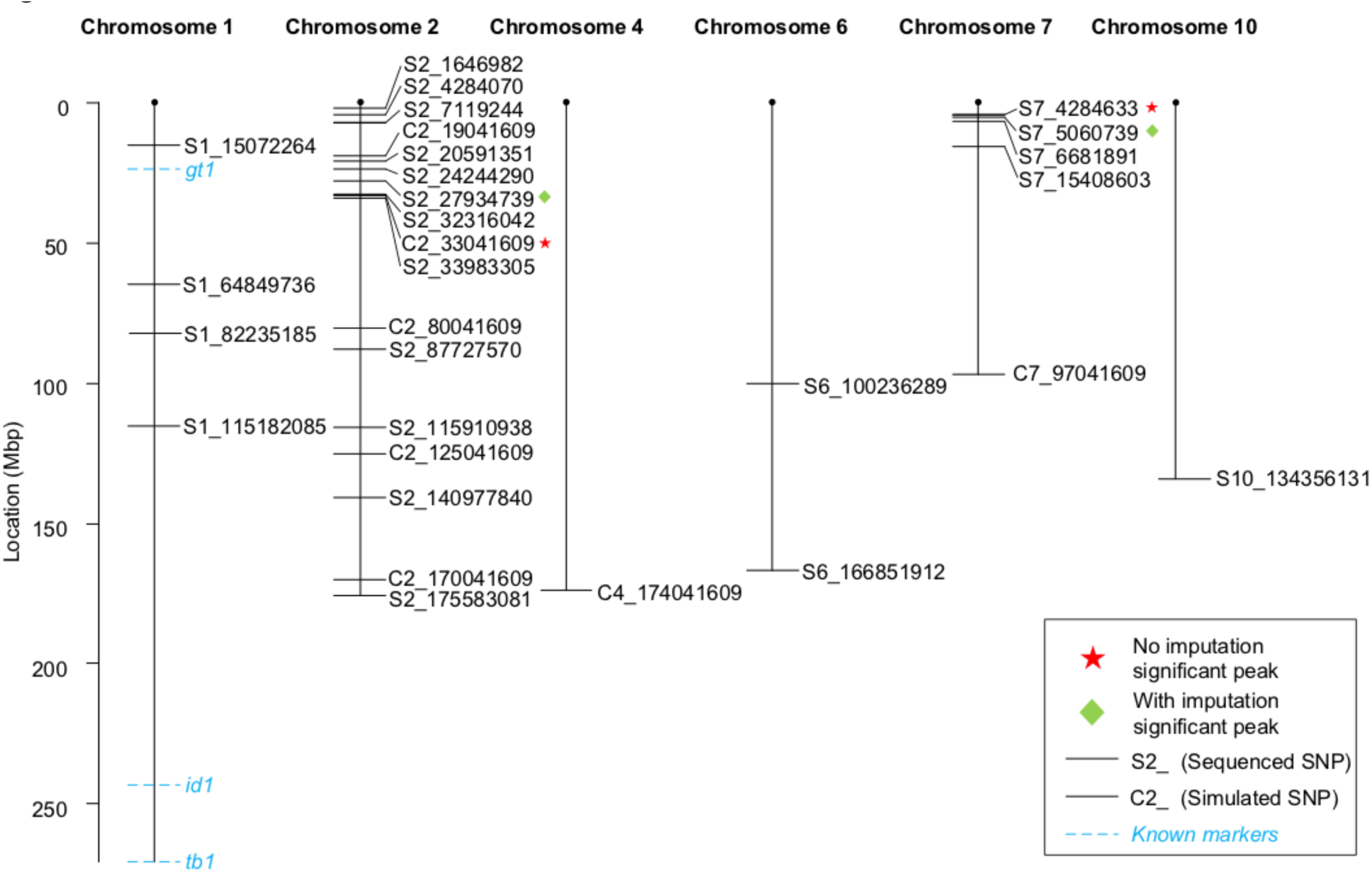
Genetic map of 30 representing SNPs and genes *gt1*, *id1*, and *tb1*. Each SNP represents a one-Mbp region except of SNP S2_27934739, which represent a SNP cluster.

In summary, the results presented here indicate that regrowth in Zea is inherited dominantly in our experimental conditions. Both the genetic and GBA analyses support a model where the regrowth trait is mainly controlled by two major regrowth loci, *reg1* and *reg2* on chromosomes 2 and 7, respectively. Even so, the data do not eliminate more complex models. Identification and functional study of the candidate genes for *reg1* and *reg2* and their possible modifiers will initiate an understanding about the molecular mechanism of perenniality in *Zea* L. We recognize that adaptability is very important for a plant to realize perennialism in a certain environment. However, this issue can be addressed separately after we understand molecularly how *Z. diploperennis* is regrowable.

## Materials and Methods

### Plant materials and phenotyping

*Zea diploperennis* (PI 462368) and *Z. mays* cv. Rhee Flint (PI 213764) were obtained from the USDA North Central Region Plant Introduction Station, Ames, IA. B73 and Mo17 inbreds were from the collection of D. Auger and are traceable back to the Maize Genetics Cooperation Stock Center, Urbana/Champaign, IL. In our designations of F_1_s and their derivatives, the female parental is shown first. All the plants used in this study were grown in the greenhouse during the winter and in the field during the summer in Brookings, SD. Controlled pollinations were done by covering tassels and ears with paper bags before and after the pollination was made. In the greenhouse, plants were maintained with a 16 h-light/8 h-dark cycle and 20/16 °C day/night temperature except that two-month old *Z. diploperennis* and its hybrid plants were treated with a 10 h light/14 h dark cycle for four weeks to induce the floral transition.

Plants were scored as regrowth if they produced shoots from the basal axillary buds after the original stalks finished flowering and senesced. Rhizome and tuber development were visually investigated on plants that were dug from the soil after senescence. TNT was investigated by counting numbers of tillers per plant when the tassel had fully emerged. Ear and kernel morphology was visually examined and photographed.

### PCR assay

DNA samples were isolated from young leaves using the CTAB procedure^(33)^ and used for PCR-based marker assay. Table 5 lists all the PCR primers used in this study. PCR assays were done using GoTaq Green Master Mix (Catalog# M7505, Promega, Madison, WI) at the following conditions: 95°C, 35 cycles of 95°C for 45 sec, 55~62°C (primer dependent, see Table 6 for detail) for 1 min and 72°C for 1 min, and 72°C for 10 min. The annealing temperatures were determined using a 1°C-touchdown PCR step starting from 65°C.

**Table 5.**
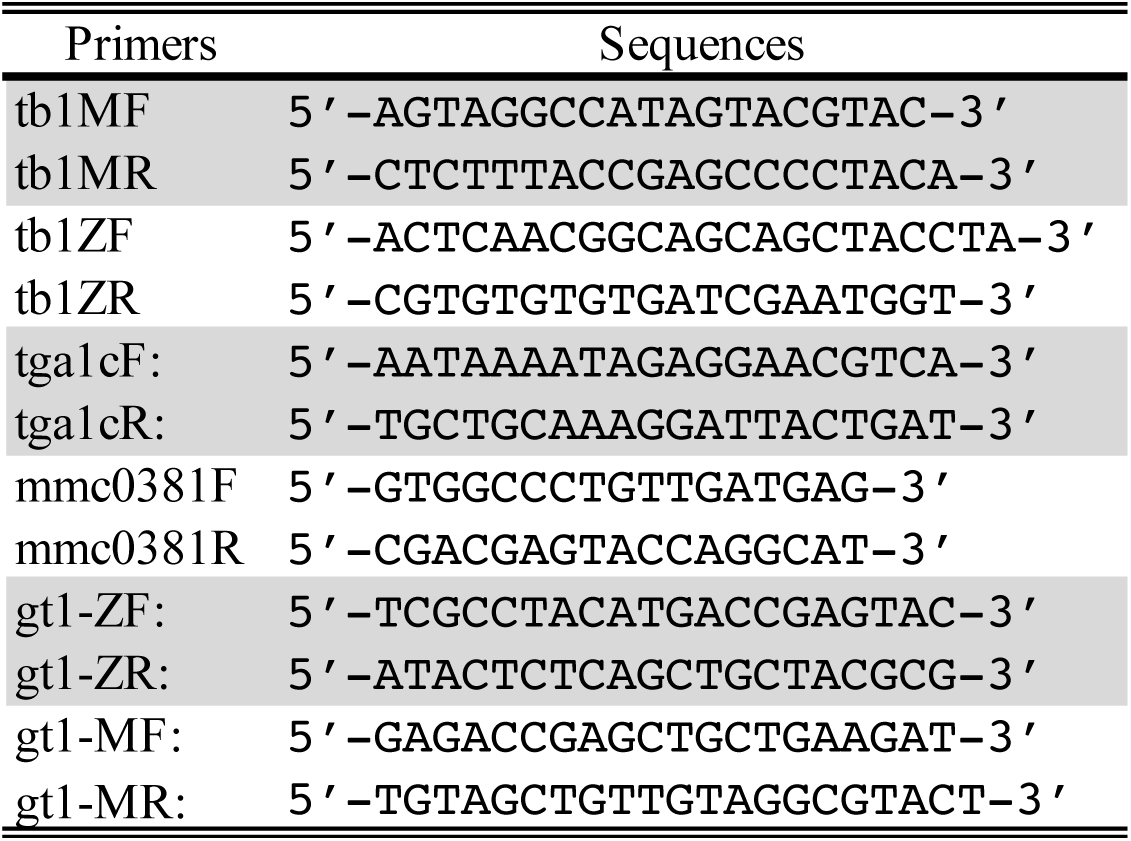
PCR primers used in this study.

### SNP discovery

The GBS assay was conducted according to Elshire and the associates^(31)^. The preparation and sequencing of the library were conducted by the University of Wisconsin Biotechnology Center (UWBRC). Generally, DNA samples were digested with *ApeK*I restriction enzyme (RE), and unique barcodes were annealed to each DNA fragments. A single-end 100 bp (1×100bp) sequencing run was carried out on an Illumina HiSeq 2500 platform. The raw data were pooled as a single fastq file and downloaded from UWBRC.

The TASSEL (Trait Analysis by Association, Evolution and Linkage) 3 pipeline was used under the guidance of TASSEL manual^(32)^ for the discovery of SNPs between *Z. diploperennis* and *Z. mays* (Supplementary Table S3). The barcoded sequence reads were collapsed into a set of unique sequence tags with counts. The tag count files were filtered for a minimum count threshold and merged into the master tag count file. The B73_RefGen_V4 reference genome sequence was downloaded from MaizeGDB and processed with Bowtie2 for alignment^(34)^. Master tags were aligned to the B73 reference genome to generate a "Tags On Physical Map" (TOPM) file, which contains the genomic position of each tag with the best unique alignment. The occupancies of tags for each taxon were observed from barcodes information in the original FASTQ files. Tag counts were stored in a "Tags by Taxa" (TBT) file. The TOPM and TBT files were used to call SNPs at the tag locations on the genome. The SNPs were filtered by minimum tag counts of 5, genotype mismatch rate of 0.1, minimum taxon coverage of 0.01, minimum site coverage of 0.2 and minimum minor allele frequency of 0.01. Fastq files containing sequences of chromosomes 1 to 10 were merged by FASTX_Toolkit and indexed. All commands for SNP discovery were executed in Ubuntu 16.04 LTS platform.

SNPs resulted from TASSEL filters plugin with a minimum minor allele frequency of 0.01 were filtered again by removing sites that had missing data in more than 20% of the B73-Zd F_2_ plants. For those SNPs that have missing data in less than 20% of the B73-Zd F_2_ plants, the missing data were imputed by treating them as heterozygote since both two alleles can be embodied and considered to be moderate. The SNPs from the 2^nd^ filtration identification were used for QTL mapping. To understand the relationship between the mapped QTL with the genetic factors revealed by the genetic analysis, the SNPs from the 2^nd^ filtration were further filtered (the 3^rd^ filtration) with χ2 (*p* < 0.05). For each regrowth-associated SNP, we expected, in the regrowable subpopulation, 33.3% plants to carry the homozygous Zd alleles (AA) and 66.6% to have the Zd-B73 heterozygous allele combination (AB) and none with the B73 homozygous alleles (BB) and, in the non-regrowable subpopulation, 14% of the plants with AA, 28.6% with AB and 57.1% with BB. Altogether, a χ2 contingency table were generated with expected 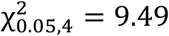. Any SNPs with χ2 < 9.49 were kept fitting the 9:7 segregation model. The 4^th^ SNP filtration was performed to keep the first SNP and remove the rest in a cluster in which all the SNPs in the range of 100 bp share the same haplotypes. In the locus analysis with the chi-square imputation, such a cluster of SNPs was treated as one locus. Removing the redundant SNPs makes the locus analysis more precise because repeated SNP sites would affect the LOD value and influence the interval estimation.

The SNPs after the 2^nd^ and the 4^th^ filtrations were used for candidate locus/QTL estimation, respectively. The locus analysis was executed by a standard QTL procedure in *R* using the *R*/qtl package (version 1.40-8)^(35)^ to better observe the contribution of each SNP and its neighbors. The *R* codes used for the analyses are listed in Supplementary Table S4. Position simulation was drawn with a maximum distance of 1.0 cM and an error probability of 1×10^−4^. The conditional genotype probability (calc.genoprob), as well as simulated genotypes (sim.geno with n.draw=32), were calculated. The “haldane” function was used to convert genetic distances into recombination fractions. Genome scan with a single locus model (scanone) was performed with a binary model using the expectation-maximization algorithm^(35)^. A permutation test with 1000 replicates was performed in scanone to visualize the LOD thresholds. We determined a locus interval by selecting the first and last SNP sites with significant LOD value. Genes within the intervals were identified by searching the corresponding region on the Gramene website.

### Statistical analyses

For statistical analyses, all genotypes and phenotypes were transformed into numeric values. For phenotypes, the regrowth plants were scored as “1” and the non-regrowth plants were scored as “2”. For genotypes, the plants that were homozygous to the *Z. diploperennis* allele were scored as “1”; those that were homozygous to the B73 or Rhee Flint allele were scored as “2”; and those that were heterozygous were scored as “3”. When conducting locus analysis, genotype “1” was transformed to “AA”, “2” to “BB” and “3” to “AB”.

A chi square goodness-of-fit test was used to find the best-fit model or linkage in the genetic analysis and reveal candidate loci on chromosomes. To determine if TNT has any correlation with regrowth, a One-Way ANOVA of TNT by regrowth was performed in JMP (JMP^®^ 11.2.0).

### Sequencing Data availability

All raw fastq data from this study are available at NCBI data deposition site (https://www.ncbi.nlm.nih.gov/bioproject/) with accession number PRJNA477673.

## Acknowledgement

This research was partially supported by funds from USDA-NIFA via South Dakota Experiment Station and Department of Biology and Microbiology, South Dakota State University. We greatly appreciate Dr. Frank M. You of Agriculture and Agri-Food Canada for his help in statistics.

## Author Contributions

Y.Y. designed and supervised this project and all the experiments, and drafted the manuscript; Y.Q., A.M, T.R., B.P., A.G., Y.Z., Y.Y. & D.A. performed the experiments and collected data; Y.Q., A.M., T.R., D.A. & Y.Y. analyzed the data; all authors discussed the results and communicated on and approved the final manuscript.

---

**Supplementary Table S1.** Segregation of regrowability among the B73-Zd F_3_s.

| **Plant** | **PT** | **Plant** | **PT** | **Plant** | **PT** | **Plant** | **PT** |
| --- | --- | --- | --- | --- | --- | --- | --- |
| BZ3-010-1-1 | 1 | BZ3-010-1-20 | 2 | BZ3-010-1-46 | 2 | BZ3-010-1-69 | 1 |
| BZ3-010-1-2 | 1 | BZ3-010-1-22 | 1 | BZ3-010-1-47 | 1 | BZ3-010-1-70 | 1 |
| BZ3-010-1-3 | 1 | BZ3-010-1-23 | 1 | BZ3-010-1-48 | 1 | BZ3-010-1-71 | 1 |
| BZ3-010-1-4 | 1 | BZ3-010-1-24 | 1 | BZ3-010-1-50 | 1 | BZ3-010-1-72 | 1 |
| BZ3-010-1-5 | 1 | BZ3-010-1-25 | 1 | BZ3-010-1-51 | 2 | BZ3-010-1-73 | 1 |
| BZ3-010-1-6 | 1 | BZ3-010-1-26 | 1 | BZ3-010-1-52 | 1 | BZ3-010-1-74 | 1 |
| BZ3-010-1-7 | 1 | BZ3-010-1-27 | 1 | BZ3-010-1-53 | 1 | BZ3-010-1-75 | 1 |
| BZ3-010-1-8 | 2 | BZ3-010-1-30 | 1 | BZ3-010-1-54 | 1 | BZ3-010-1-76 | 2 |
| BZ3-010-1-9 | 2 | BZ3-010-1-31 | 1 | BZ3-010-1-56 | 1 | BZ3-010-1-77 | 2 |
| BZ3-010-1-11 | 2 | BZ3-010-1-32 | 1 | BZ3-010-1-57 | 1 | BZ3-010-1-78 | 1 |
| BZ3-010-1-12 | 1 | BZ3-010-1-33 | 2 | BZ3-010-1-68 | 2 | BZ3-010-1-79 | 1 |
| BZ3-010-1-13 | 2 | BZ3-010-1-34 | 1 | BZ3-010-1-59 | 1 | BZ3-010-1-80 | 1 |
| BZ3-010-1-14 | 2 | BZ3-010-1-35 | 1 | BZ3-010-1-61 | 2 | BZ3-010-1-81 | 2 |
| BZ3-010-1-15 | 1 | BZ3-010-1-38 | 2 | BZ3-010-1-62 | 1 | BZ3-010-1-82 | 2 |
| BZ3-010-1-16 | 1 | BZ3-010-1-39 | 1 | BZ3-010-1-63 | 1 | BZ3-010-1-83 | 2 |
| BZ3-010-1-17 | 1 | BZ3-010-1-40 | 1 | BZ3-010-1-64 | 2 | BZ3-010-1-84 | 1 |
| BZ3-010-1-18 | 1 | BZ3-010-1-41 | 2 | BZ3-010-1-67 | 1 | BZ3-010-1-85 | 1 |
| BZ3-010-1-19 | 1 | BZ3-010-1-42 | 2 | BZ3-010-1-68 | 1 | BZ3-010-1-87 | 1 |

1: regrowth; 2: non-regrowth

**Supplementary Table S2.** Phenotypes and the *gt1*, *id1* and *tb1* haplotypes of 26 F_2_ plants and three F_3_ populations of the *Zea mays* cv. Rhee Flint x *Z. diploperennis* cross.

| Line | PT | *tb1* | *id1* | *gt1* | Line | PT | *tb1* | *id1* | *gt1* | Line | PT | *tb1* | *id1* | *gt1* |
| --- | --- | --- | --- | --- | --- | --- | --- | --- | --- | --- | --- | --- | --- | --- |
| F<sub>2</sub>-1 | 1 | 3 | 1 | 3 | F<sub>3</sub>-3-2 | 1 | 1 | 2 | 1 | F<sub>3</sub>-5-13 | 1 | 1 | 2 | 1 |
| F<sub>2</sub>-2 | 2 | 2 | 2 | 2 | F<sub>3</sub>-3-3 | 1 | 1 | 2 | 1 | F<sub>3</sub>-5-14 | 2 | 1 | 2 | 1 |
| F<sub>2</sub>-3 | 1 | 1 | 2 | 3 | F<sub>3</sub>-3-4 | 2 | 1 | 2 | 3 | F<sub>3</sub>-5-15 | 2 | 1 | 2 | 1 |
| F<sub>2</sub>-4 | 2 | 2 | 2 | 2 | F<sub>3</sub>-3-6 | 1 | 1 | 2 | 3 | F<sub>3</sub>-5-16 | 2 | 1 | 2 | 1 |
| F<sub>2</sub>-5 | 1 | 1 | 2 | 1 | F<sub>3</sub>-3-7 | 1 | 1 | 2 | 1 | F<sub>3</sub>-9-1 | 1 | 1 | 2 | 1 |
| F<sub>2</sub>-6 | 1 | 3 | 1 | 1 | F<sub>3</sub>-3-8 | 1 | 1 | 2 | 3 | F<sub>3</sub>-9-2 | 1 | 1 | 2 | 2 |
| F<sub>2</sub>-7 | 2 | 2 | 2 | 2 | F<sub>3</sub>-3-9 | 1 | 1 | 2 | 3 | F<sub>3</sub>-9-3 | 1 | 1 | 2 | 2 |
| F<sub>2</sub>-9 | 1 | 1 | 2 | 3 | F<sub>3</sub>-3-10 | 1 | 1 | 2 | 3 | F<sub>3</sub>-9-4 | 1 | 1 | 2 | 3 |
| F<sub>2</sub>-10 | 2 | 3 | 1 | 3 | F<sub>3</sub>-3-11 | 2 | 1 | 2 | 3 | F<sub>3</sub>-9-5 | 1 | 1 | 2 | 1 |
| F<sub>2</sub>-11 | 1 | 2 | 2 | 3 | F<sub>3</sub>-3-12 | 1 | 1 | 2 | 1 | F<sub>3</sub>-9-6 | 2 | 1 | 2 | 2 |
| F<sub>2</sub>-12 | 1 | 3 | 1 | 1 | F<sub>3</sub>-3-13 | 1 | 1 | 2 | 3 | F<sub>3</sub>-9-7 | 1 | 1 | 2 | 3 |
| F<sub>2</sub>-13 | 2 | 3 | 1 | 3 | F<sub>3</sub>-3-14 | 1 | 1 | 2 | 1 | F<sub>3</sub>-9-8 | 1 | 1 | 2 | 1 |
| F<sub>2</sub>-14 | 2 | 2 | 2 | 3 | F<sub>3</sub>-3-15 | 1 | 1 | 2 | 1 | F<sub>3</sub>-9-9 | 1 | 1 | 2 | 3 |
| F<sub>2</sub>-15 | 1 | 3 | 1 | 1 | F<sub>3</sub>-3-16 | 2 | 1 | 2 | 2 | F<sub>3</sub>-9-10 | 1 | 1 | 2 | 3 |
| F<sub>2</sub>-16 | 2 | 2 | 2 | 1 | F<sub>3</sub>-5-1 | 1 | 1 | 2 | 1 | F<sub>3</sub>-9-11 | 1 | 1 | 2 | 1 |
| F<sub>2</sub>-17 | 1 | 3 | 1 | 3 | F<sub>3</sub>-5-2 | 1 | 1 | 2 | 1 | F<sub>3</sub>-9-12 | 1 | 1 | 2 | 3 |
| F<sub>2</sub>-18 | 2 | 3 | 1 | 2 | F<sub>3</sub>-5-3 | 2 | 1 | 2 | 1 | F<sub>3</sub>-9-13 | 1 | 1 | 2 | 1 |
| F<sub>2</sub>-19 | 2 | 3 | 1 | 2 | F<sub>3</sub>-5-4 | 2 | 1 | 2 | 1 | F<sub>3</sub>-9-14 | 2 | 1 | 2 | 3 |
| F<sub>2</sub>-20 | 1 | 3 | 1 | 3 | F<sub>3</sub>-5-5 | 1 | 1 | 2 | 1 | F<sub>3</sub>-9-15 | 2 | 1 | 2 | 3 |
| F<sub>2</sub>-21 | 2 | 2 | 2 | 3 | F<sub>3</sub>-5-6 | 1 | 1 | 2 | 1 | F<sub>3</sub>-9-16 | 1 | 1 | 2 | 1 |
| F<sub>2</sub>-22 | 2 | 2 | 2 | 3 | F<sub>3</sub>-5-7 | 1 | 1 | 2 | 1 | F<sub>3</sub>-12-1 | 1 | 1 | 2 | 1 |
| F<sub>2</sub>-23 | 1 | 2 | 2 | 3 | F<sub>3</sub>-5-8 | 1 | 1 | 2 | 1 | F<sub>3</sub>-12-2 | 1 | 1 | 2 | 1 |
| F<sub>2</sub>-24 | 1 | 1 | 2 | 3 | F<sub>3</sub>-5-9 | 1 | 1 | 2 | 1 | F<sub>3</sub>-12-3 | 1 | 1 | 2 | 1 |
| F<sub>2</sub>-25 | 1 | 2 | 1 | 1 | F<sub>3</sub>-5-10 | 1 | 1 | 2 | 1 | F<sub>3</sub>-12-4 | 1 | 3 | 1 | 1 |
| F<sub>2</sub>-26 | 1 | 3 | 2 | 1 | F<sub>3</sub>-5-11 | 2 | 1 | 2 | 1 | F<sub>3</sub>-12-5 | 1 | 3 | 1 | 1 |
| F<sub>3</sub>-3-1 | 1 | 1 | 2 | 3 | F<sub>3</sub>-5-12 | 2 | 1 | 2 | 1 |  |  |  |  |  |

\* For phenotype (PT), "1" is for regrowth and "2" is for non-regrowth; for genotypes, "1" is homozygous for the *Zea diploperennis* allele, "2" is homozygous for the *Z. mays*'s allele and "3" is for heterozygous.

**Supplementary Table S3.**
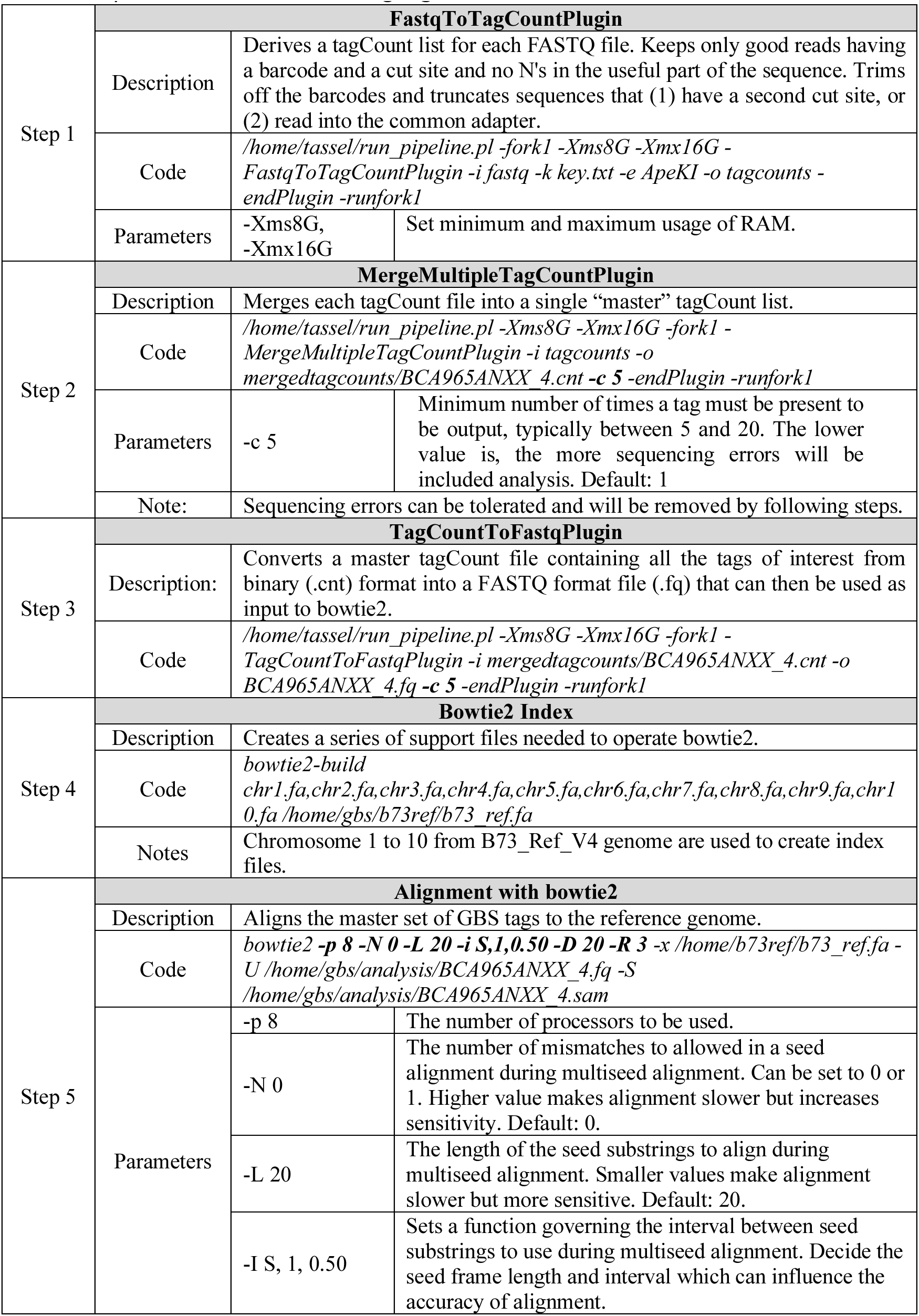
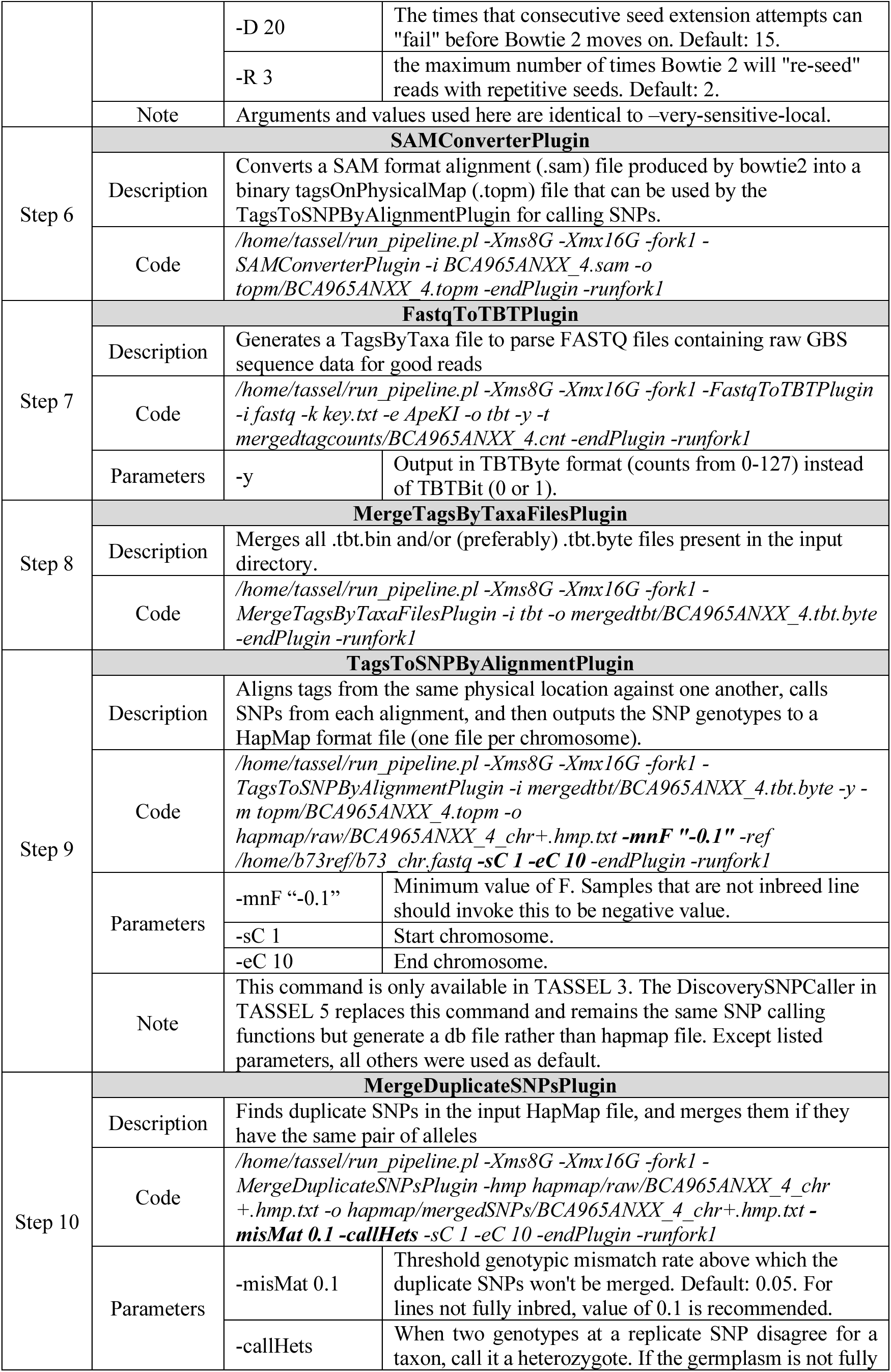
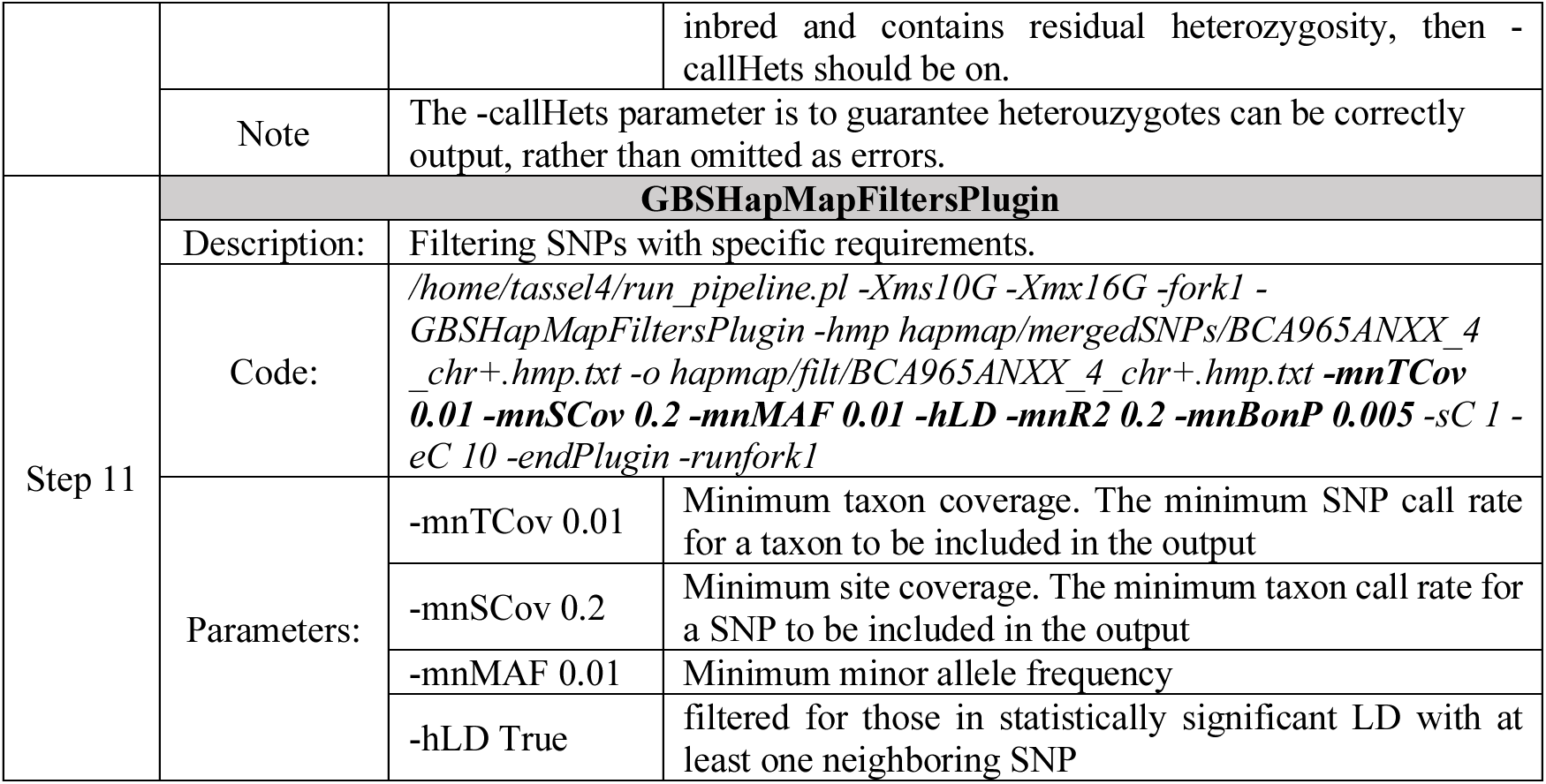
Steps and codes in TASSEL pipeline including example command lines and brief descriptions. Parameters are highlighted and described when first be used.

**Supplementary Table S4.**
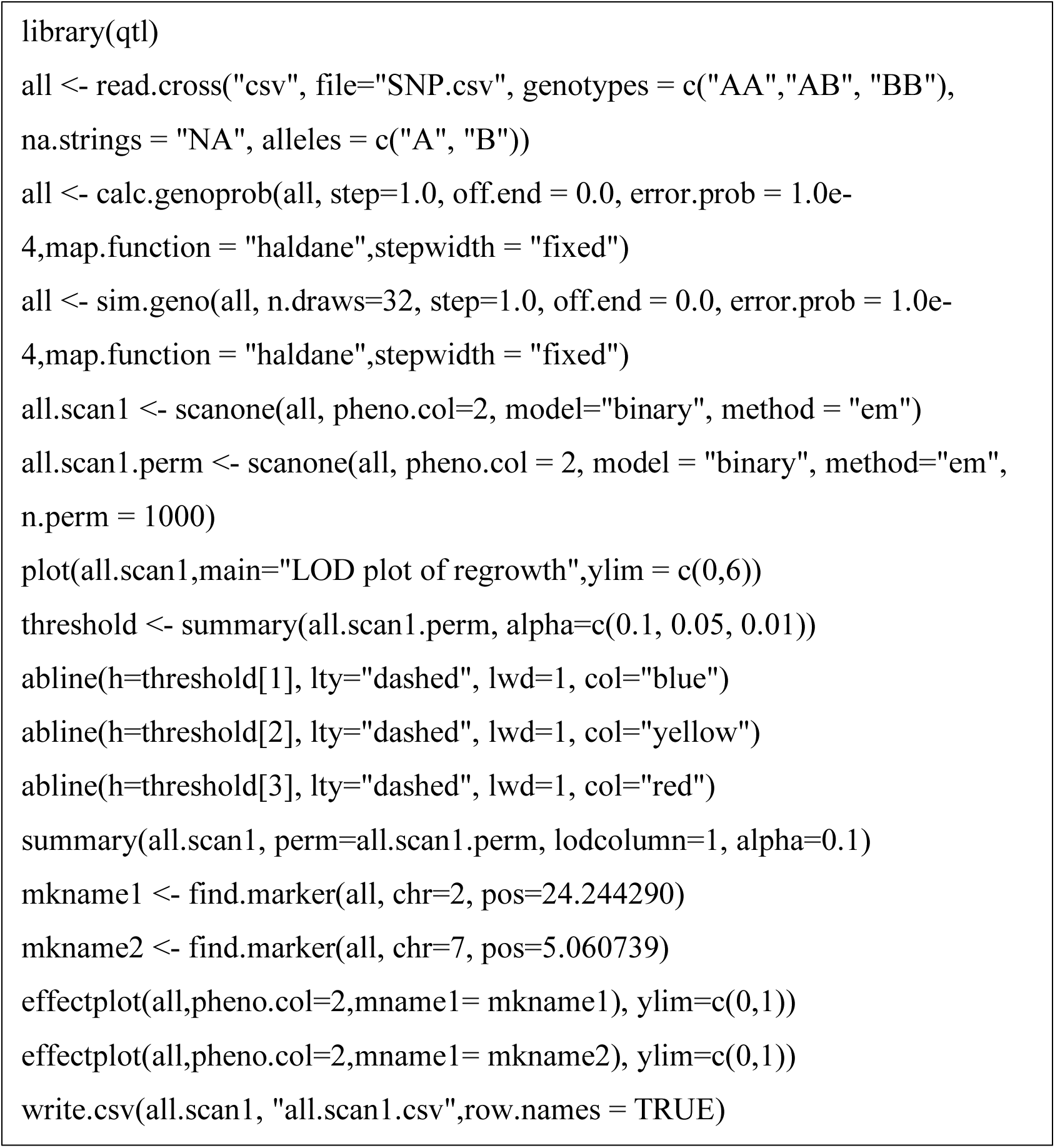
R codes used for candidate locus/QTL analyses.

**Supplementary Figure S1.**
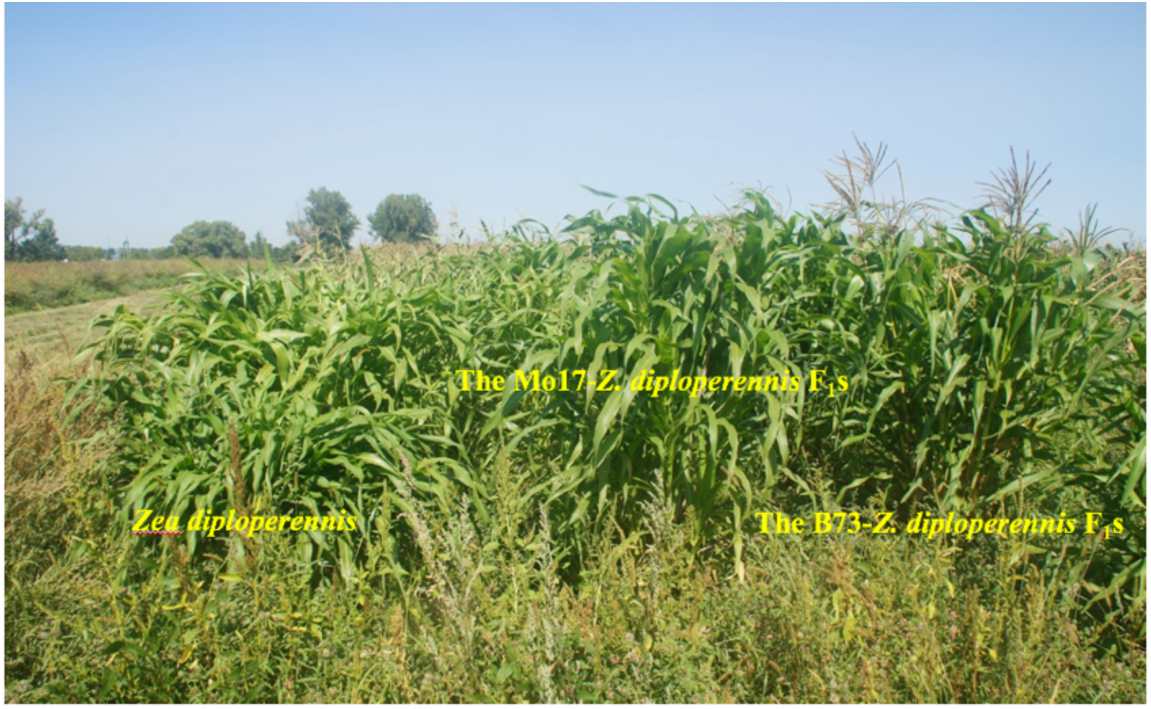
A photo showing the growth of *Zea diploperennis* and its F_1_ with Z. mays B73 or Mo17 in the field in the Summer 2017.

**Supplementary Figure S2.**
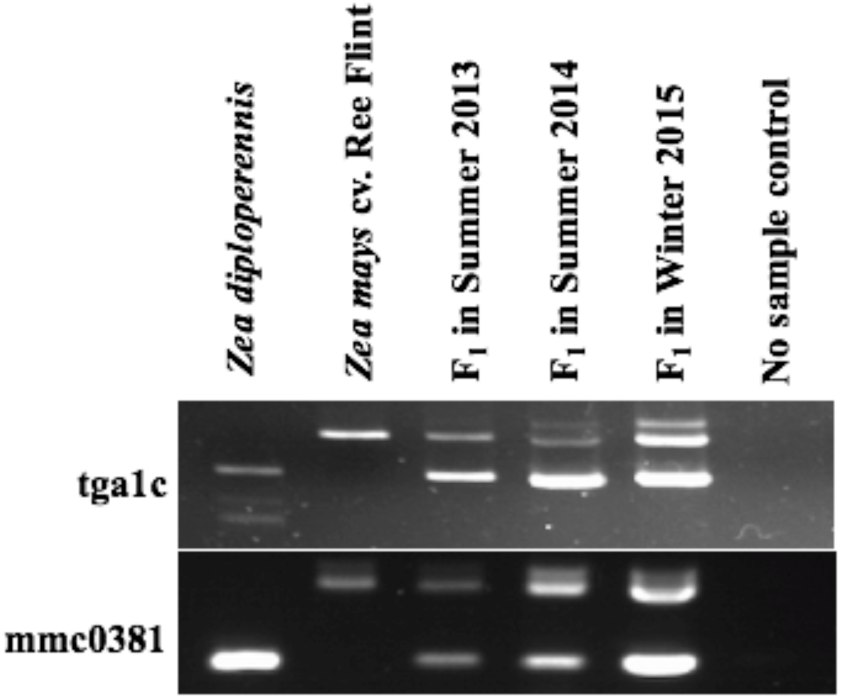
An agarose gel image showing that two molecular markers confirmed the heterozygosity of a *Z. diploperennis*-*Z. mays* cv. Rhee Flint F_1_ plant over three life-cycles.

**Supplementary Figure S3.**
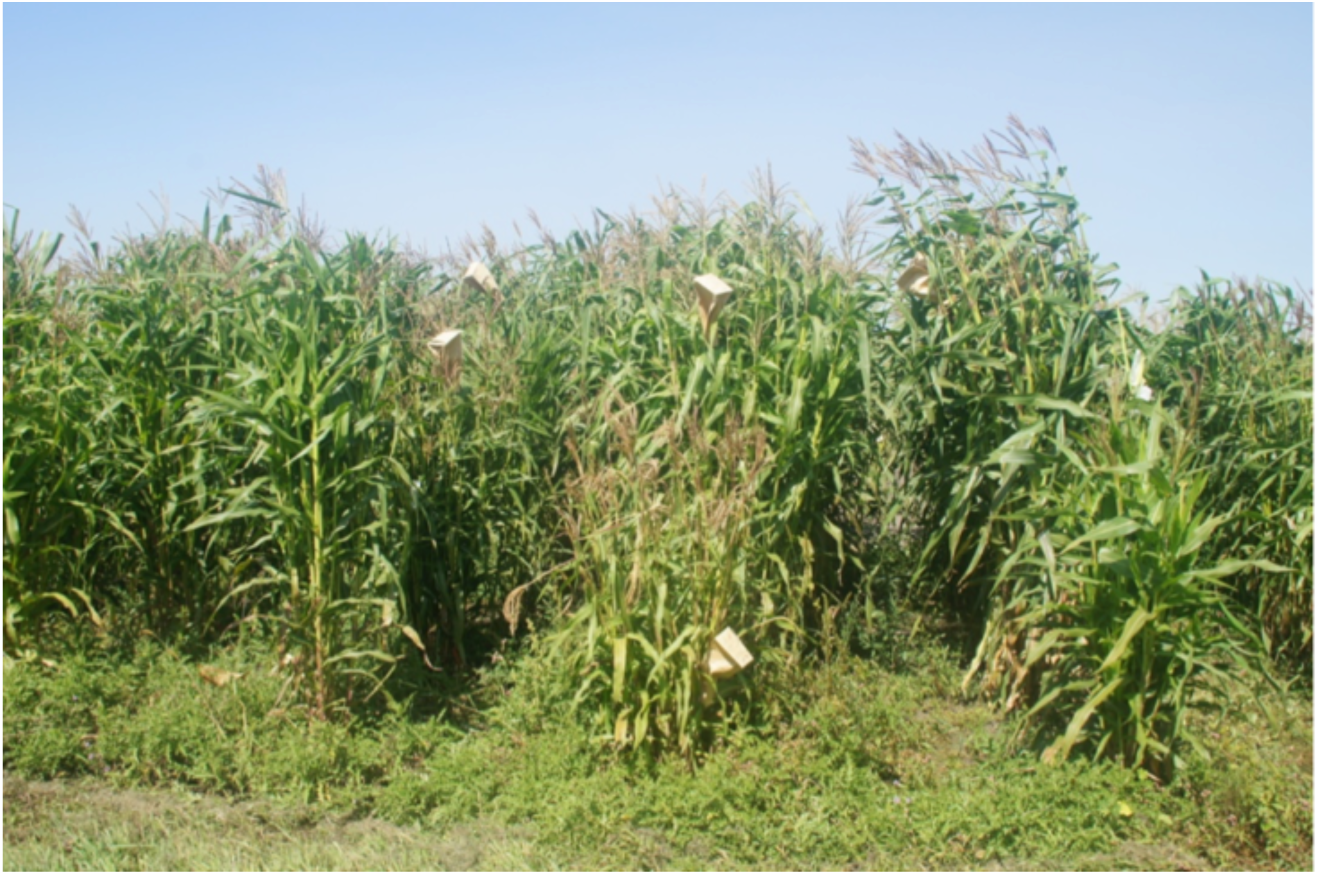
A photo showing the regrowth of the B73 - *Z. diploperennis* F4s in the summer, 2017.

